# Local Adaptation to the Sex Environment: Reciprocal Sex-Limited Selection in Different Thermal Regimes

**DOI:** 10.64898/2026.01.23.701411

**Authors:** Chloe Melo-Gavin, Michelle J. Liu, Aneil F. Agrawal

## Abstract

The shared genome prevents each sex from independently responding to the selection experienced by that sex. We used experimental evolution in *Drosophila melanogaster* with separate pools of Chromosome 3s for males (male-limited chromosomes) and females (female-limited chromosomes) for 15 generations. Viewing each sex as a separate environment, we performed a reciprocal transplant between the sexes to quantify the strength of local adaptation to each sex environment. Each chromosome type was more beneficial in the sex it had been selected for (i.e., local adaptation to sex). Because it has been postulated that sex differences in selection may depend on how well adapted a population is to the abiotic environment, we performed experimental evolution at two thermal regimes: one benign temperature to which the populations were well-adapted and one novel temperature. Female-specific adaptation was stronger at the benign temperature whereas male-specific adaptation was stronger in the novel temperature. Within chromosome pools, male and female fitness were more positively correlated in the novel compared to the benign temperature. Though males carrying male-limited chromosomes were typically more fit than males carrying female-limited chromosomes, they were also more harmful to their female mating partners.

## Introduction

Sex differences in selection on autosomal variants, in either magnitude or direction, will cause each sex to evolve differently than it would if it could evolve independently of the other sex. We might expect that each sex is typically hindered by sharing a gene pool with the other because the response to selection on the alleles most important to one sex can be diluted by weaker, or even opposing selection, in the other sex. However, if selection on autosomal variants is typically concordant but on average stronger in one sex than the other, then the sex experiencing weaker selection may adapt faster by sharing a gene pool with the more strongly selected sex. We are still at the early stages of understanding how much sharing a genome affects the adaptation of each sex. Are both sexes equally affected by sharing a genome? Are the sexes affected more strongly by sharing a genome under some conditions than others?

Answers to such questions depend on the frequency of variants that experience sex-differential selection and on the type of differential selection. This would require us, ideally, to know how much of the genetic variance in fitness is due to (i) sexually antagonistic alleles, (ii) alleles with sex limited effects (i.e., alleles affecting fitness in one sex but not the other), and (iii) alleles that are selected in the same direction in both sexes, but with different magnitudes. As it is not currently possible to address this issue directly at a genome-wide level, one approach is through the detailed study of specific alleles and their fitness effects in each sex (e.g., Sharp and Agrawal, 2008; Melde et al., 2024; Barson et al., 2015; Hawkes et al., 2016). However, only a small number of variants can be studied this way and the variants chosen for any given study may not be representative of natural variation more broadly.

Alternatively, one can examine the net effect of variation across many loci. These approaches provide only limited insight into the details of the underlying genetic architecture but more directly address higher level questions regarding sex-specific adaptation with a shared genome. One such approach is the measurement of the intersexual genetic covariance in fitness (Koch et al., 2022; Delcourt et al., 2012; Foerster et al., 2007; Svensson et al., 2009). This covariance can be used—in principle—to predict how adaptation in one sex is hindered or helped by sharing a gene pool with the other. Still, the match between predicted and realized responses to selection in quantitative genetics tends to become progressively worse across more than a handful of generations (Roff, 2012; Hansen et al., 2011).

Another approach, and the primary focus in our work, is to use experimental evolution to more directly examine the consequences of selection in the absence of a shared gene pool. A classic example of this is the study of Prasad et al. (2007) who performed experimental evolution in *Drosophila melanogaster,* using a method first pioneered by Rice (1996,1998), to create a separate pool of haplotypes that exclusively experienced father to son transmission and thus only experienced selection in males (i.e., male-limited selection). After 25 generations of this selection, male-limited haplotypes improved male fitness compared to controls but decreased female fitness and additionally led to more masculinized phenotypes for both sexes (Prasad et al., 2007; Abbott et al., 2010). Similarly, both Abbott et al. (2020) and Grieshop et al. (2025) performed male-limited selection using different methods and also found that male-limited chromosomes improved male (but not female) mating success. Rice (1992) saw chromosomes with female-limited regions improved female fitness (and decreased male fitness) and Lund-Hansen et al. (2020) observed phenotypic feminization in response to their female-limited selection. In the hermaphroditic worm *Macrostomum lignano*, when selection was limited to only male- or female-function, the experimental populations appeared to increase their fitness through the selected-sex function relative to the alternatively selected populations (Nordén et al., 2023). Across these studies, these improvements in male-specific or female-specific fitness provide evidence that there were sex-specific adaptations that had been hindered under normal inheritance.

The experiment here was inspired by these past experiments but differs from most of them in two important ways. First, we implemented a reciprocal design (see also Nordén et al., 2023 and Morrow et al., 2008), creating two pools of genetically variable Chromosome 3 and enforcing a pattern of same-sex inheritance such that one pool was inherited from mother to daughter, and experienced selection only in females (i.e., Female-Limited (“FL”) selection) and the other pool was passed down father to son and experienced selection only in males (i.e., Male-Limited (“ML”) selection). This reciprocal design allows us to quantify sex-specific adaptation by re-purposing the reciprocal transplant framework commonly used for local adaptation (e.g., quantifying female-specific adaptation by measuring how much more fit FL than ML chromosomes are in the “female environment”). We first ask if there is divergence in the fitness effects of ML and FL lines, as expected if selection differs between the two sexes. We then compare the strength of local adaptation to each sex. A difference implies that the sexes are affected differently by sharing a genome. For example, if we find adaptation to the female environment is strong (females with FL chromosomes have vastly superior fitness compared to females with ML chromosomes), but adaptation to the male environment is weak (males with ML or FL chromosomes have similar fitness), this would imply that, under normal inheritance female fitness is affected by sharing a genome more than male fitness.

A novel aspect of our study is that we perform the experimental evolution in two temperature environments: the standard lab temperature and a novel thermal regime. Cryptic genetic variation is expected to be revealed (or magnified) in novel environments (Paaby and Rockman 2014). Most cryptic variation is likely to be sexually concordant whereas standing variation in a constant environment will be comparatively enriched for sexually antagonistic variation (Long et al., 2012; Connallon and Hall, 2016). In addition, changes in the environment may re-align the direction of selection between the sexes, causing the effects of alleles to be less sexually antagonistic than in more stable environments (Connallon and Hall, 2016). Consistent with this, de Lisle et al.’s (2018) meta-analysis examined 722 paired estimates of selection across 25 species and found that male and female selection coefficients were more similar when species were in less stable environments or far from the centre of their range (i.e., presumably where populations are least well adapted).

In our experiment, one might expect the existence of variants that were neutral in the original temperature but are adaptive (or deleterious) for both sexes in the novel thermal regime. If these alleles are more strongly selected in males than females because of the alignment of sexual and natural selection (Whitlock and Agrawal, 2009; Rowe and Rundle, 2021), then in the novel thermal regime, more so than the standard temperature, one might expect stronger local adaptation of ML chromosomes to males than the local adaptation of FL chromosomes to females (that is, males with ML chromosomes would have superior fitness compared to males with FL chromosomes, but females with FL chromosomes would have similar fitness to females with ML chromosomes). Thus, we examine how local adaptation to “sex”–and its asymmetry between the sexes–varies between populations evolving at the standard temperature and those evolving in the novel thermal regime to gain insight into how the effect of the shared genome varies across environments.

In addition to comparisons between FL and ML pools, we also examine variation within these pools. Specifically, we examine the intersexual correlation for fitness, *r_MF_*, and test whether this correlation is more positive at the novel temperature than at the benign temperature, as predicted if novel environments reveal sexually concordant variation (or cause the alignment of previously antagonistic variants). Some studies find evidence of more negative *r_MF_*in well-adapted compared to novel environments (Long et al., 2012; Han and Dingemanse, 2017; Berger et al., 2014), but results are mixed (Delcourt et al., 2012; Koch et al., 2020; Punzalan et al., 2014; Martinossi-Allibert et al., 2018).

Finally, the male fitness assays provided the opportunity to examine how sex-limited selection affected the evolution of mate harm. In *Drosophila,* both pre- and post-mating male traits can give males a competitive advantage while reducing the fitness of their female mating partner (e.g., aggressive courtship, harmful seminal fluid proteins; Patridge et al., 1986; Nandy et al., 2013; Chapman et al., 1995). This mate harm can be mediated, at least in part, via traits present in both sexes (e.g., body size, locomotor activity; Pitnick and García–González, 2002; Nandy et al., 2013) that are known to be differentially selected for between the sexes (Long et al., 2007; Abbott et al., 2010). Even though our experimental regime does not change selection on mate harm directly, separating the gene pool of males and females would allow divergence in the frequency of alleles affecting mate harm if such alleles were previously constrained by the shared gene pool.

## Methods

### Base Population

The source population used for experimental evolution was initially created by mixing more than 25 inbred lines of *Drosophila melanogaster* from multiple sources and allowing them to recombine for many generations to generate a diverse set of haplotypes. By the time the experiment began most lines had been recombining for more than 110 generations and all lines had been recombining for over 60 generations. As part of this process, two recessive phenotypic markers on Chromosome 3, *se^1^* (located on 66D5 of Chromosome 3R) and *hh^bar3^* (Cytological position 94E1 on Chromosome 3R) were introgressed into the population to aid in sorting flies during experimental evolution. That is, during experimental evolution, individuals that were homozygous for either marker were discarded (see below). The process of introgression occurred in two stages. First more than 25 inbred lines were mixed and the *se^1^* marker was introgressed into this population. The version of this population containing the *se^1^*marker (but not the *hh^bar3^* marker) we refer to as the “Grand Ancestor” population. Afterwards, the *hh^bar3^* marker was introgressed into a copy of the Grand Ancestor population to produce what we refer to as the “Ancestor” population (though the Grand Ancestor population was also maintained). The end result of this multi-year preliminary phase was the Ancestor population that carried *se^1^* and *hh^bar3^* but was genetically diverse elsewhere on Chromosome 3. The other chromosomes (Chromosomes 2, 4, *X* and *Y*) in the Ancestor population were also genetically variable but are not the focus here. A detailed description of the history of the source population is provided in the Supplementary Material.

This Ancestor population homozygous for *se^1^* and *hh^bar3^* was used, along with two *TM6B* balancer stocks, to establish the experimental populations. One *TM6B* balancers was marked with the dominant phenotypic bristle marker *Sb* and contained a construct with a dominant fluorescent marker, *DsRed* (the full genotype of the construct, *DsRed Hr5.IE1 OpIE2-neoR traF OpIE2-pacR dsx* , is described in Kandul et al. (2019)). We hereafter referred to this balancer as *TM6B DsRed Sb.* The other balancer was only marked with the *Sb* marker and is referred to as *TM6B Sb. TM6B Sb* and *TM6B DsRed Sb* are recessive lethal. Both balancer stocks were backcrossed to the Ancestor 2-4 times prior to their use, with the final backcrosses involving ∼1200 Ancestor flies for each stock. (Note, the non-balancer 3^rd^ chromosomes in these two resulting stocks were exclusively from Ancestor population).

From these two stock populations, ∼1000 *TM6B DsRed Sb/se^1^ hh^bar3^* females were collected and mated to ∼1000 *TM6B Sb/se^1^ hh^bar3^* males in 13 bottles (40mL standard yeast media, ∼150 flies per bottle) to begin Generation 0 of the experimental populations. This cross was flipped into new bottles 5 times (over 12 days), resulting in 6 sets of 13 bottles, one for each experimental population. The offspring from this cross were considered Generation 1 and sorted following our experimental design described below.

### Experimental Evolution Protocol

For 15 generations, Chromosome 3 (∼40% of the genome) gene pools of males and females were separated as follows. In our experimental populations, we had two “pools” of genetically variable copies of Chromosome 3 (hereafter referred to as “focal” chromosome pools) that experienced selection, and two pools of alternative balancer chromosomes (*TM6B Sb* and *TM6B DsRed Sb*) that functioned to keep the focal chromosome pools separated from one another (**FIGURE 1).**

**FIGURE 1:**
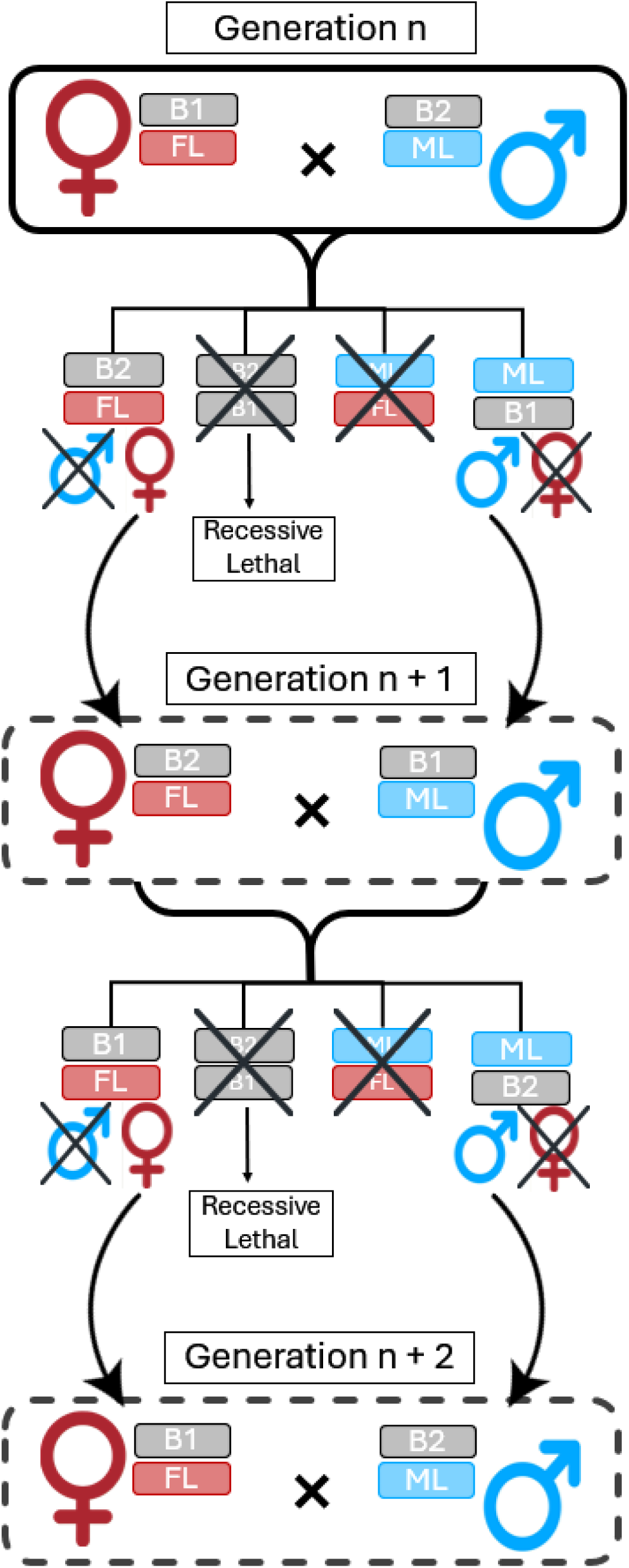
Sex-limited experimental evolution protocol. We performed 15 generations of sex-limited experimental evolution in *Drosophila melanogaster* at both a benign temperature and in a novel thermal regime. We created two pools of genetically variable Chromosome 3, and enforced a pattern of same-sex inheritance such that one pool was inherited from mother to daughter, and experienced selection only in females (i.e., Female-Limited (“FL”) selection) and the other pool was passed down father to son and experienced selection only in males (i.e., Male-Limited (“ML”) selection). The cross design, which repeats every two generations, is shown.

Each generation we selected for flies that inherited a focal chromosome from their same-sex parent and a balancer from their different-sex parent. Practically, this can be achieved because all potential mothers carried one type of balancer, and all potential fathers carried the other type of balancer (and the two balancer types are phenotypically distinguishable from each other). By manually sorting the two balancer types into alternate sexes each generation, one pool of focal chromosomes was passed down from mother to daughter and experienced selection only in females (i.e., Female-Limited (“FL”) selection) and the other pool was passed down father to son and experienced selection only in males (i.e., Male-Limited (“ML”) selection) (**FIGURE 1**). In this way each pool of chromosomes was free to increase the frequency of beneficial alleles in that sex (ex. female beneficial alleles for FL pool) regardless of the effect in the other sex (ex. males). This separation of gene pools thereby removed any constraint of the shared genome for these chromosomes and allowed sex-specific adaptation that would have been prevented under normal inheritance.

This sex-limited selection occurred at two treatment temperatures (3 populations per treatment, *N* = 1050), one benign temperature treatment (25° C) to which the populations are well-adapted, and one novel temperature treatment (mostly at 28° C, see below). All 6 populations were maintained on a three-week cycle (12:12 hour light:dark cycle at 50% relative humidity) on regular yeast-sugar media. Flies were mixed in small cages on Day 14 of each generation, then sorted (as described above **FIGURE 1**) into interaction vials (∼40 vials of 13 males paired with 13 females for each population). After seven days of interaction, flies were transferred to oviposition vials to lay eggs for 24-28 hours, after which adults were removed, beginning Day 1 of the next generation cycle. For the first two days of each generation, individuals in the novel temperature treatment were kept at 18°C to synchronize generation time between all temperature treatments (i.e., all populations on the same three-week cycle).

### 2 by 2 Competitive Fitness assays

After 15 generations of experimental evolution, ∼ 30 individual ML and ∼ 30 individual FL chromosomes per population were isolated into individual lines with the ancestral background and the balancer *TM6B Sb* (**FIGURE S2**). Competitive fitness was measured for each ML and FL line in both sexes using individuals which were heterozygotes for a focal chromosome and *TM6B Sb*, matching one of the genetic backgrounds in which evolution of the focal chromosomes had occurred. We housed 6 focal individuals of a given sex with 6 same-sex, fluorescent competitors and 12 different-sex Grand Ancestors. Grand Ancestors flies were chosen instead of Ancestors because Ancestor flies are homozygous for *hh^bar3^* resulting in significantly impaired eye development and viability compared to wildtype, though *hh^bar3^* heterozygotes (like the focal flies) are not significantly different from wildtype in this regard (Rogers et al., 2005). In the female fitness assays, the female competitors had the genotype *w-; DsRed Hr5.IE1 OpIE2-neoR traF OpIE2-pacR dsx* (Kandul et al., 2019), which contains the dominant fluorescent marker, *DsRed*. In the male fitness assays, the male competitors were from another stock homozygous for *DsRed*. Mimicking the maintenance described above, females and males were combined on Day 14 of their generation cycle and given seven days of interaction, after which females were moved into oviposition vials for 24 hours. For the female fitness assay, focal and competitor females were housed in the same oviposition vial. For the male fitness assay, females were split into two separate oviposition vials. Focal flies were assayed at their respective treatment temperature (25° C for the benign temperature 28° C for the novel temperature). All female deaths (competitor and focal females in the female fitness assay or the females in the male fitness assay) were recorded up until oviposition time. In both male and female assays, competitive fitness was calculated as the proportion of (non-fluorescent) offspring produced by the focal flies as measured on Day 14 after oviposition.

The female fitness assay was conducted in 2 experimental blocks, and the male fitness assay was conducted in 3 blocks. Grand Ancestor competitive fitness was also measured in each block (i.e., Grand Ancestors were the ‘focal’ flies in these replicates). When measuring Grand Ancestor competitive fitness, flies were collected from the Grand Ancestor population on Day 14 of their maintenance cycle, and 6 randomly chosen *se^1^* homozygous individuals of the same sex were grouped together per each replicate vial. Note that in assays of ML or FL chromosomes, the focal flies carried a balancer chromosome as they did during experimental evolution (see above). In contrast, in vials where Grand Ancestors were the focal flies, the focal flies did not carry a balancer. We use these Grand Ancestor assays as a standard for normalization to facilitate combining data across blocks rather than as a representation of fitness effects at the onset of sex-limited evolution.

For some analyses, we calculated fitness of ML and FL lines relative to the Grand Ancestor (hereafter ‘relative fitness’) as the fitness per line per sex, divided by the mean Grand Ancestor fitness for each Temperature/Sex/Block combination. The bootstrapping done for **FIGURE 2** involved first randomly sampling the raw counts of both individual focal lines and Grand Ancestor replicates, after which we calculated fitness of the sampled lines relative to the mean of the sampled Grand Ancestor replicates.

**FIGURE 2:**
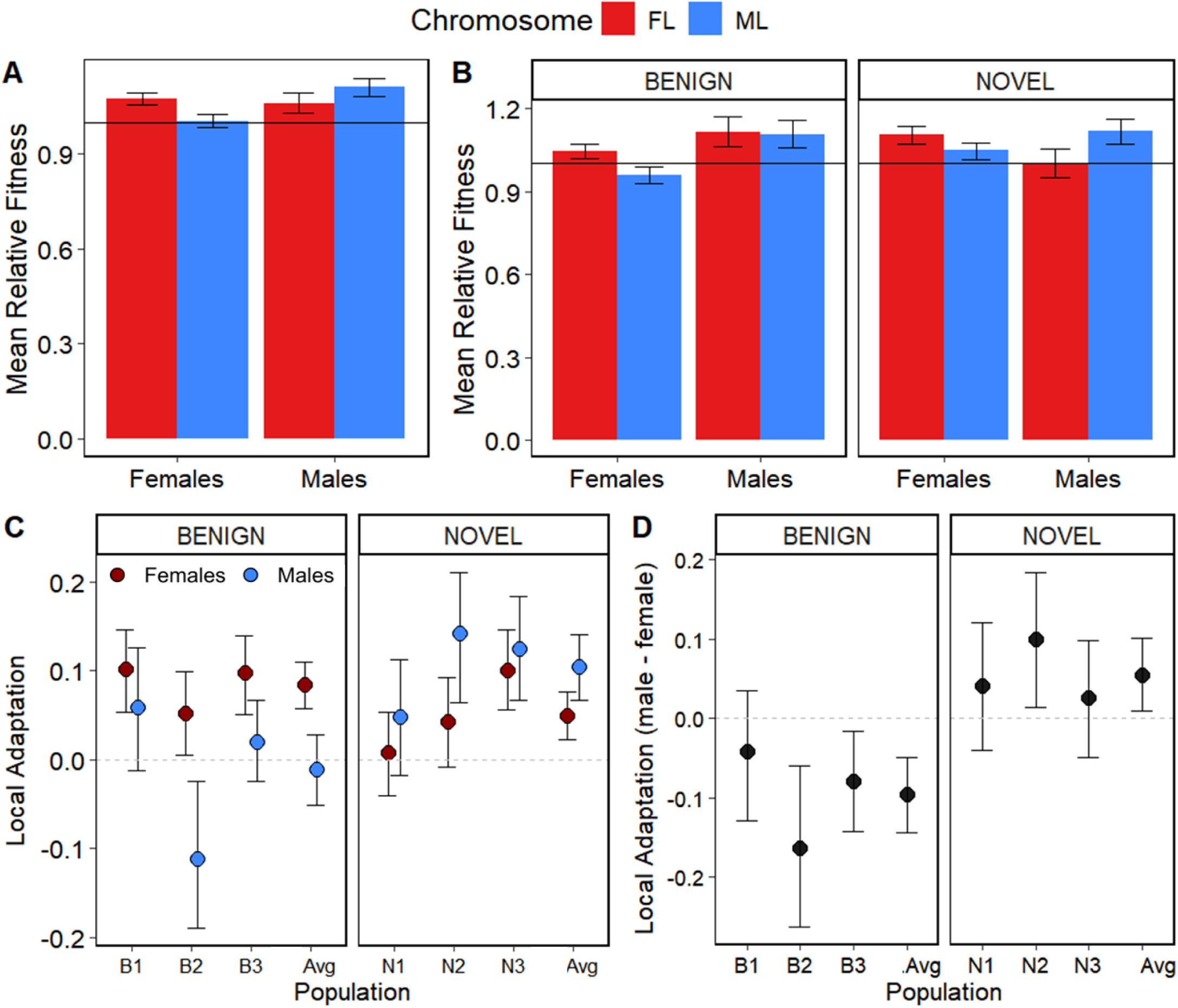
Relative fitness of ML and FL chromosomes in both sexes. (A) Comparison of relative fitness between chromosome types, averaging over both treatment temperatures. Fitness is shown relative to the Grand Ancestor (black line). (B) Comparison of fitness in the benign and novel thermal treatments. (C) Local adaptation of FL and ML chromosome pools to the female and male environments, respectively. (D) Sex difference in local adaptation (male - female). Error bars are 95% bootstrapped confidence intervals of the mean.

### Divergence between ML and FL

Variation in competitive fitness was examined using a quasi-binomial linear mixed model using the *lme4* package in *R* (Bates et al., 2015). Sex (i.e., male or female fitness assay), Chromosome Pool (ML or FL), and Treatment Temperature were considered as fixed effects while Block, Population, and Replicate were considered random effects. Because of a significant Sex by Chromosome by Temperature interaction, we also analyzed each temperature separately.

### Quantifying Local Adaptation

For each population, we quantified how well each pool was adapted to being in its respective sex. We compared relative fitness using a “local vs foreigner” approach (Kawecki and Erbert, 2004). Female-specific adaptation (local adaptation to the female environment) for population *P* was measured as: 𝐿𝐴_𝐹,𝑃_ = 𝑤̅_𝐹,𝐹𝐿,𝑃_ − 𝑤̅_𝐹,𝑀𝐿,𝑃_/(𝑚𝑎𝑥 (𝑤̅_𝐹,𝐹𝐿,𝑃_, 𝑤̅_𝐹,𝑀𝐿,𝑃_) where 𝑤̅_𝑆,𝐶,𝑃_is the average relative fitness of individuals of sex *S* constructed with chromosome type *C* from population *P*. Conversely, male-specific adaptation (local adaptation to the male environment), was measured as 𝐿𝐴_𝑀,𝑃_ = 𝑤̅_𝑀,𝑀𝐿,𝑃_ − 𝑤̅_𝑀,𝐹𝐿,𝑃_/(𝑚𝑎𝑥 (𝑤̅_𝑀,𝐹𝐿,𝑃_, 𝑤̅_𝑀,𝑀𝐿,𝑃_). We performed 1000 bootstraps for each chromosome type in each population by resampling relative fitness values in each population and recalculating local adaptation for each sex per sample. For an individual population, sex-specific adaptation was said to be significant if the confidence intervals did not overlap 0.

For each population, we evaluated the difference in the strength of female-specific adaptation and male-specific adaptation as Δ𝐿𝐴_𝑃_ = 𝐿𝐴_𝑀,𝑃_ − 𝐿𝐴_𝐹,𝑃_. This was done for all 1000 bootstrap estimates to determine bootstrap 95% confidence intervals. Local adaptation was considered stronger in one sex if the confidence intervals did not overlap 0. Separately for each of the two temperature treatments, we also measured the average local adaptation to each sex across the three populations (𝐿̅̅𝐴̅̅_𝑆_ = 𝐿𝐴_𝑆,𝑃1_ + 𝐿𝐴_𝑆,𝑃2_ + 𝐿𝐴_𝑆,𝑃3_)/3) as well as the sex difference in local adaptation (Δ̅̅𝐿̅𝐴̅̅ = Δ𝐿𝐴_𝑃1_ + Δ𝐿𝐴_𝑃2_ + Δ𝐿𝐴_𝑃3_)/3), using bootstrapping to determine bootstrap 95% confidence intervals.

### Intersexual Correlation for Fitness r_MF_

We examined the correlation between female fitness and male fitness, *rMF*, at the line level, across populations, as follows. For each sex (*S*) separately, we centered the relative fitness of each line with respect to the mean relative fitness for each Chromosome type (*C*) per each Population (*P*) 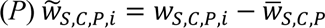. We then calculated the Pearson correlation between centered male fitness and centered female fitness for each Chromosome type within each temperature treatment (novel temperature ML, novel temperature FL, benign temperature ML and benign temperature FL).

We performed 1000 bootstraps for each chromosome type by resampling lines with their centered male and female fitness values and calculating the Pearson correlation coefficient (estimate of *r_MF_*) for each sample. To examine differences in *r_MF_* across temperature treatments, we subtracted the benign *r_MF_* estimate from the novel *r_MF_* estimate, separately for each chromosome pool; this was done for all 1000 bootstrap estimates to determine bootstrap 95% confidence intervals on the difference between temperature treatments for each chromosome pool. If the 95% CI did not overlap 0, then the pools were considered significantly different.

Lastly, to compare the temperature treatments holistically, we averaged *r_MF_* estimates from the FL and ML pools for each temperature treatment and then took the difference between these temperature-specific averages (difference = novel – benign). Bootstrap confidence intervals of this difference were determined as above.

We emphasize that the intersexual correlation estimated here is not the intersexual *genetic* correlation as would be measured in a quantitative genetic study designed with that purpose. Here we have only a single male and female replicate vial for each line and thus cannot partition genetic and environmental contributions to the correlation. However, as male and female fitness measures for each line were done in separate blocks using individuals from different developmental vials, environmental effects will be negligible on the intersexual covariance but still contribute to the variance within each sex. Consequently, one would expect the *r_MF_* we estimate is biased downwards in magnitude relative to the true intersexual genetic correlation.

It should be noted that the genetic differences among lines are not solely due to the focal (ML or FL) chromosomes. Although all lines were constructed using the same procedure, we expect some genetic differences among lines on other chromosomes due to segregating variation in the stocks used in line construction. However, this source of “extraneous” genetic variation should be equivalent within each ML or FL set of lines from each population (i.e., there should be no difference, on average, in background between ML and FL lines). Though this extraneous genetic variation will contribute to each of our estimates of *r_MF_*, the focal chromosomes are expected to be the primary source of differences in *r_MF_*among different sets of lines.

### Mate harm

The male fitness assay provided us with the opportunity to examine how males carrying ML versus FL chromosomes affected the females with which they were housed. For each line in the male fitness assay, we examined female mortality (the number of female deaths during mating) and female fecundity (total offspring produced by females surviving to the oviposition period). Analysis of female mortality in the male fitness assay was performed using a quasi-binomial linear mixed model with Chromosome Pool and Treatment Temperature as main effects and Block, Population and Replicate as random effects. Female fecundity was analyzed using a gaussian linear mixed model with Chromosome Pool, Number of Female Deaths, and Treatment Temperature as main effects and Block and Population as random effects. Number of Female Deaths (or, equivalently, Number of Surviving Females) was included to examine variation in vial productivity controlling for the number of females surviving to the oviposition period.

We also examined the phenotypic correlation between male fitness and fitness of females with whom they were housed in the male fitness assays. For male fitness, we used centered male relative fitness values described above. For female fitness, we used total offspring produced by females per each focal male line, which we then centered for each chromosome type per population. We then calculated the Pearson correlation between centered male fitness and centered female fitness for each chromosome type within each temperature treatment. Note that fitness of focal males is a competitive measure (proportion of offspring sired by focal males) whereas the fitness of the associated females is an absolute measure (total offspring produced in a vial) and the two measures will have some autocorrelation as the latter number is in the denominator of the former. To account for this, we generated a null distribution by randomizing the pairing between focal offspring number and competitor offspring number (for a given chromosome pool, within a given population and block) and recalculating male relative fitness and female fitness from the sum of these randomized pairings and then generating the correlation for each chromosome pool within each temperature treatment. We performed this 1000 times to generate a list of 1000 correlation coefficients for each of the four chromosome pools. The observed correlation was determined to be significantly different if it lay outside the 95% confidence intervals of the null distribution.

## Results

### Competitive Fitness

We created separate gene pools that experienced sex-limited selection. One pool of chromosomes was passed down from mother to daughter and experienced selection only in females (i.e., Female-Limited (“FL”) selection) and the other pool was passed down father to son and experienced selection only in males (i.e., Male-Limited (“ML”) selection). This separation of gene pools should have allowed each gene pool to increase the frequency of alleles beneficial within the corresponding sex regardless of the effect in the other sex. However, if selection is the same for both sexes, Female-Limited and Male-Limited chromosomes should have the same effects on fitness as one another in both sexes.

Consistent with sex differences in selection, there was a significant Chromosome by Sex effect on competitive fitness (*p* < 0.05; **FIGURE 2A**; see **Table 1** for full model results; see also **SUPPLEMENTAL FIGURE 4**), where females carrying FL chromosomes (hereafter “FL-females”) had higher fitness than females carrying ML chromosomes and males had higher fitness when carrying ML chromosomes compared to FL chromosomes. This Chromosome by Sex effect persisted when Treatment temperatures were examined individually (Chromosome by Sex effect: *p <* 0.01 for both benign and novel temperatures) but there were notable differences between Treatment temperatures (Chromosome by Sex by Treatment temperature effect: *p* < 0.05; **FIGURE 2B**).

**Table 1:** Analysis of Competitive Fitness. A comparison of Focal (Non-Fluorescent) compared to Competitor (Fluorescent) offspring using a quasi-binomial linear mixed model. Significant effects are bolded.

| | $\chi^2$ | DF | P-value |
| --- | --- | --- | --- |
| **Chromosome** | **6.10** | **1** | **0.0136** |
| Sex | 0.869 | 1 | 0.351 |
| Treatment | 3.44 | 1 | 0.0638 |
| **Chromosome × Sex** | **5.44** | **1** | **0.0196** |
| Chromosome × Treatment | 0.0178 | 1 | 0.893 |
| **Sex × Treatment** | **6.01** | **1** | **0.0142** |
| **Chromosome × Sex × Treatment** | **5.70** | **1** | **0.0169** |

We then quantified local adaptation to each sex environment for each population, using a “local vs foreign” comparison of fitness. Consistent with the linear model, all six FL chromosome pools showed positive local adaptation to females (i.e., female-specific adaptation), four of which were significant (i.e., confidence intervals do not overlap 0; **FIGURE 2C**). The mean female-specific adaptation across the three replicate populations was significantly positive for each of the temperature treatments. Five of the six ML chromosome pools showed positive male-specific adaptation, two of which were individually significantly positive. The mean male-specific adaptation across the three replicate populations at the novel temperature was significantly positive but the mean for the benign temperature populations had confidence intervals that overlapped 0.

We then compared the strength of local adaptation to each sex, within each population (i.e., the difference in male- vs. female-specific adaptation, Δ𝐿𝐴 = 𝐿𝐴_𝑀_ − 𝐿𝐴_𝐹_). The point estimate for Δ𝐿𝐴 was negative (i.e., stronger female- than male-specific adaptation) in all three benign temperature populations, and significantly so in two of them (**FIGURE 2D**). The mean of Δ𝐿𝐴 across these three populations was significantly negative. In contrast, the point estimate for Δ𝐿𝐴 was positive (i.e., stronger male- than female-specific adaptation) in all three novel temperature populations, though only significantly so in one. The mean of Δ𝐿𝐴 across these three populations was significantly positive.

### Intersexual correlation for fitness r_MF_

We next examined the intersexual correlation for fitness (*r_MF_*) within chromosome pools. We centered relative fitness values for each chromosome/sex/population combination to remove population differences and measure a single *r_MF_* for ML and FL pools across populations for each temperature treatment. In the benign temperature , both FL and ML chromosome pools had an *r_MF_* indistinguishable from 0 (Pearson correlation: *r_MF(ML)_* = - 0.007, *p* = 0.945; *r_MF(FL)_* = 0.002, *p* = 0.843; **FIGURE 3**). In contrast, in the novel temperature, ML lines and FL lines had a significantly positive *r_MF_*(*r_MF(ML)_* = 0.266, *p* < 0.001 and *r_MF(FL)_* = 0.240, *p* < 0.01 respectively). For both the ML and FL chromosomes, novel temperature lines had significantly higher *r_MF_* than benign temperature lines (*r_MF(Novel_ML)_ - r_MF(Benign_ML)_* = 0.281. 95% bootstrapped confidence interval: [0.043, 0.442] and *r_MF(Novel_FL)_ - r_MF(Benign_FL)_* = 0.202 [ 0.039, 0.435]). To compare the temperature treatments holistically, we averaged *r_MF_* estimates from ML lines and from the FL lines within a temperature treatment. Averaging across the ML and FL, the novel temperature treatment had a significantly higher *r_MF_* than the benign temperature (*r_MF(Novel_avg)_ - r_MF(Benign_avg)_* =

**FIGURE 3:**
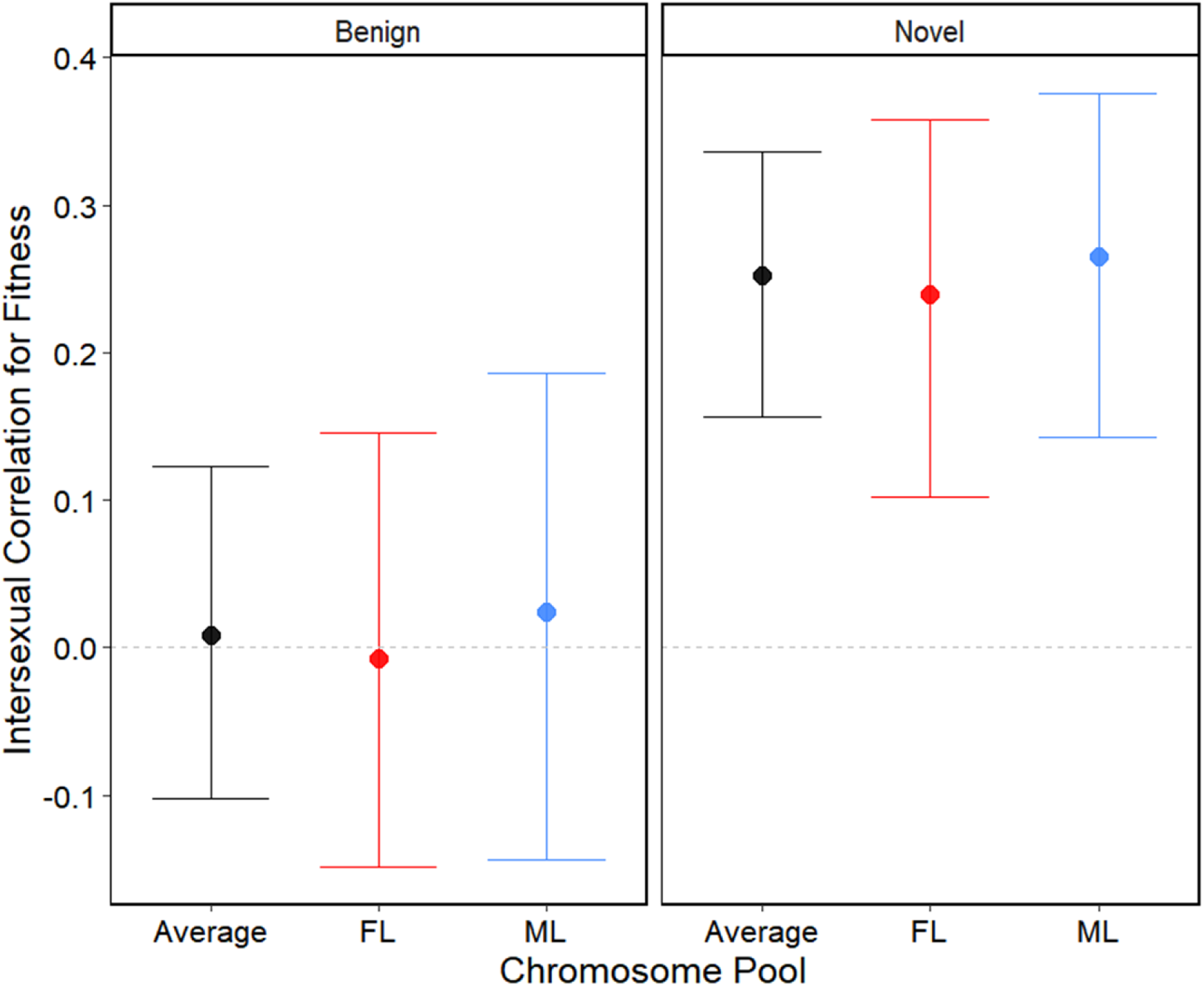
Pearson correlation of male and female relative fitness per line in both the benign and novel thermal treatments. Error bars represent 95^%^ bootstrapped confidence intervals.

0.234 [ 0.105, 0.385]).

### Mate Harm

The male fitness assay provided the opportunity to examine how removing the shared genome affected male’s harmfulness to their female mating partners. We examined the fecundity and mortality of females present in the male fitness assays, comparing between females housed with males possessing ML or FL chromosomes. Because only half the males in each assay vial were focal males (the remainder were competitor males, common across all replicates), this difference underestimates the true divergence in harm between ML-males and FL-males. With respect to the analysis of female survival, there is a significant temperature treatment by chromosome effect (Treatment by Chromosome effect; *p* < 0.05), reflecting the higher mortality of females housed with ML-males (rather than FL-males) in the novel but not benign temperature treatment (**FIGURE 4A**). Of those females that survived, females housed with ML-males had significantly fewer total offspring than those housed with FL-males (Chromosome effect *p* < 0.001; **FIGURE 4B**). This divergence in mate harm between chromosome pools suggests that sex-specific selection on mate harm is constrained under normal inheritance.

**FIGURE 4:**
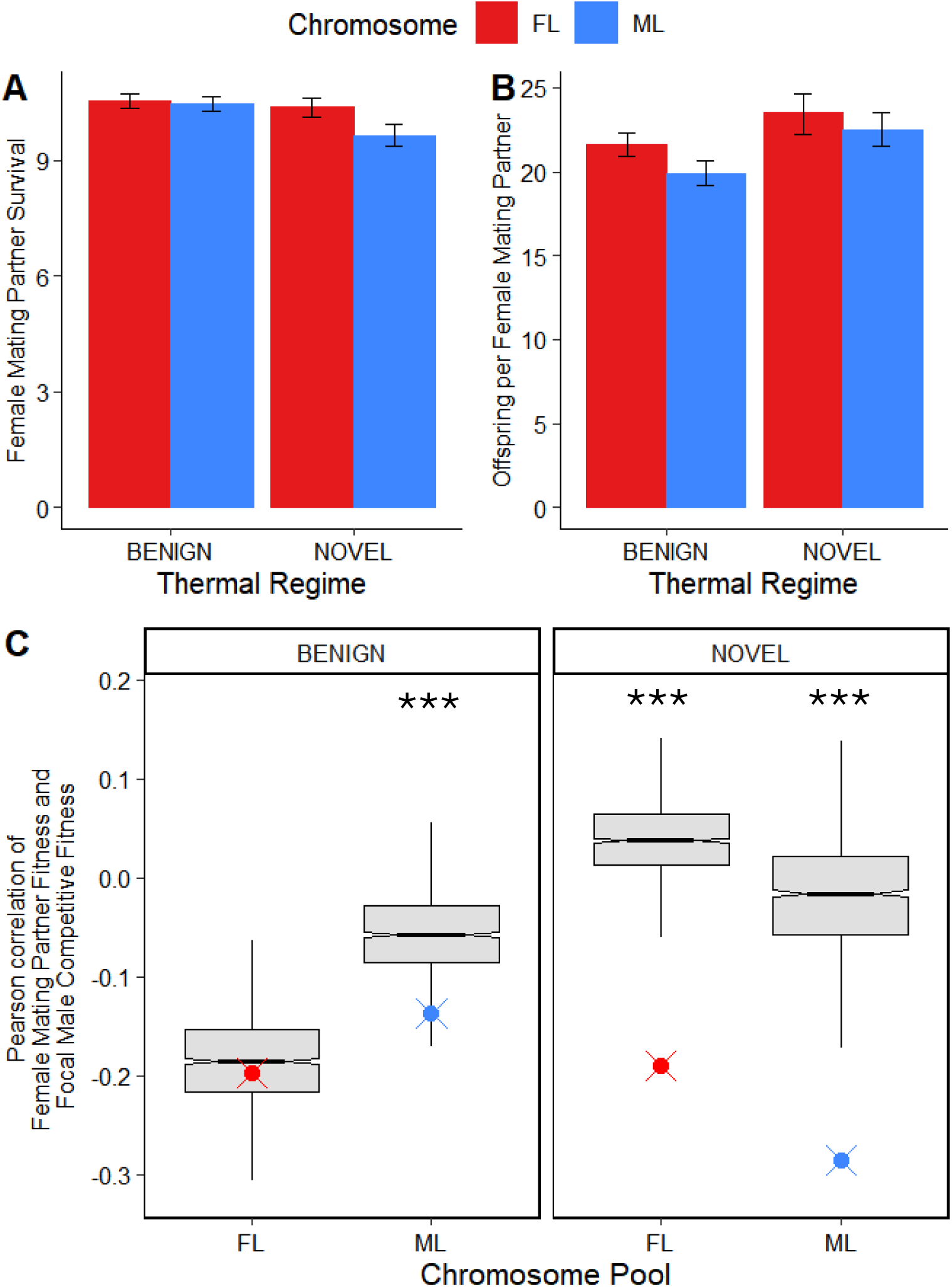
Male-induced harm on females. (A) Survivorship of females housed with a male carrying either a ML or FL chromosome (B) Total offspring produced per surviving female when housed with a male carrying either a ML or FL chromosome. Error bars are 95% confidence intervals of the means. Note the statistical analysis reported in the text was instead based on modeling the number of offspring per vial including the number of surviving females as a covariate. Either representation gives qualitatively the same pattern. (C) Points represent observed Pearson correlation of focal male competitive fitness and their respective female mating partner fitness (total offspring produced) compared to a null distribution (boxplot). Asterisks indicate significance, i.e., the observed correlation is unlikely to come from the null distribution (*p* < .05).

Additionally, within chromosome pools, we found a negative correlation between male relative competitive fitness (proportion of offspring sired by focal males) and the number of offspring produced by females with whom they were housed (here using total offspring produced per replicate vial, thereby integrating male effects on both female survival and fecundity). This suggests that male genotypes which are more harmful also had higher competitive fitness (**FIGURE 4C).**

## Discussion

We expected that creating separate gene pools for males and females would cause these pools to diverge in a sex-specific manner as each pool adapted to its respective sex, a prediction which was largely fulfilled in both thermal regimes. This is consistent with previous sex-limited experiments that have found sex-limited selection led to an increase in fitness for the selected sex (Martinossi-Allibert et al., 2019; Grieshop et al., 2025; Morrow, et al., 2008; Prasad et al., 2007; Rice, 1998; Rice, 1992; Nordén et al., 2023; Abbott et al., 2020), suggesting sex-specific fitness was hindered under normal inheritance (i.e., when sharing a genome). The fitness differences between females (or males) carrying FL- or ML-chromosomes imply that sex-specific fitness is hindered by the shared genome at both well-adapted and novel temperatures. Sex-independent selection should not contribute to differences between ML and FL chromosomes, meaning that local adaptation to each sex should result from the response to sex differences in selection (in either direction or magnitude) that would have been constrained or diluted when sharing a genome. That FL and ML chromosomes displayed sex-dependent fitness effects and local adaptation to their sex environment suggests there are sex differences in selection in both temperature treatments.

Our reciprocal design allowed us to quantify sex-specific adaptation by measuring how much more fit FL (ML) chromosomes are than ML (FL) chromosomes in the female (male) environment. From the differences in the strength of local adaptation to each sex, we can infer the sexes are not affected equally by sharing a gene pool. Local adaptation to females was stronger than to males in the benign (i.e., standard laboratory) thermal regime whereas local adaptation to males was stronger in the novel thermal regime (**FIGURE 2**). These asymmetries in the strength of local adaptation suggest that female fitness is more affected by sharing a genome than male fitness at the benign temperature whereas the reverse is true in the novel thermal regime.

One potential explanation for the latter result may be that if novel environments tend to increase the amount of sexually concordant variation (Connallon and Hall, 2014), then, because variants typically experience stronger selection in males than females (Whitlock & Agrawal 2009, Janicke et al., 2016), this would lead to a faster increase of concordantly beneficial alleles in the ML pool compared to the FL pool. Consequently, ML chromosomes would perform reasonably well in females, while the “lag” in the FL pool would cause the reverse consequence for FL chromosomes in males, resulting in lower female than male local adaptation.

The asymmetry in local adaptation is a direct assessment of the asymmetry in how each sex is affected by the gene pool being selected through the other sex rather than its own. However, the asymmetry in local adaptation can be somewhat misleading with respect to inferences about how much each sex is constrained from achieving its optimal genotype when evolving under a shared gene pool. As a toy example, consider sexually antagonistic alleles at mutation-selection balance. Imagine that in the ancestral (benign) environment, some low frequency variants are strongly deleterious in males but weakly beneficial in females. Such alleles would increase in frequency in the FL pool, resulting in the appearance of stronger male than female local adaptation (because FL chromosomes make very unfit males). It would be wrong to conclude from the asymmetry in local adaptation that, with a shared gene pool, males were more strongly constrained than females from reaching their optimum. To directly address the issue, one would need populations in which the gene pool was shared between the sexes and compared to ML and FL pools. Due to the logistical demands of maintaining these evolving populations, we were unable to keep such populations. Nonetheless, our experiment provides clear evidence that the sexes are affected differently by selection in the other sex and that this pattern reverses between the benign and novel thermal regimes.

Though most other related studies have imposed sex-limited selection in only one sex, we are aware of two that attempted to impose sex-limited selection in both (Morrow et al 2008, Norden et al. 2023). In neither case was the explicit goal of the study to directly examine sex differences in how each sex was affected by selection in the other, and for both studies there are reasons (described below) to have concerns about making this comparison. Nonetheless, from visual inspection of their reported results, both studies found evidence for slightly weaker female than male local adaptation, which could be viewed as opposite to our results.

Working with *Drosophila melanogaster*, Morrow et al. (2008) attempted to limit selection to only one sex at a time for 26 generations. Selection was prevented in the other sex by experimentally preventing any variance in fitness in that sex. Doing so required maintaining male- and female-limited populations under different conditions from one another and this could easily have resulted in the two treatments not being symmetrical removals of selection on the other sex. For example, in the treatment intending to (only) eliminate selection on males, flies were maintained under monogamy (single male-female pair) conditions. Such conditions also alter selection on females (Long et al. 2009; Yun et al. 2018) and likely change, rather than simply eliminate, selection on males (Holland and Rice 1999; Crudgington et al. 2005, Yun et al. 2021).

Nordén et al. (2023) performed experimental evolution on the hermaphroditic worm *Macrostomum lignano*. For 14 generations, a *GFP* marker was experimentally forced to be inherited exclusively through only the male- or female-function. The region linked to the marker thus experiences sex-limited selection. A nice feature of that experiment was the inclusion of control populations in which the marker was inherited equally through both sexes. They found significant sex-dependent fitness effects in the populations undergoing female-limited selection but not in male-limited populations, which performed similarly to the control for both male and female fitness. Changes in gene expression compared to control populations where also much larger in the female-limited populations than in the male-limited populations (Cīrulis et al., 2023). One possible reason male-limited populations performed similarly to control populations could be that the ancestor populations were already closer to the male fitness optima than the female fitness optima (i.e., female fitness was more constrained). Alternatively, as suggested by the authors, the experimental evolution limited the opportunity for selection on male function by limiting the number of possible mates. Another possible unintended asymmetry between female- and male-limited selection treatments concerns the region linked to the *GFP* marker. If recombination rates differ between male and female meiosis, as is common (Sardell and Kirkpatrick, 2020), then the two treatments will differ in how much of the genome experiences sex-limited selection, making it difficult to compare them. An additional complication in comparing studies of sex-limited selection in hermaphrodites to those in species with separate sexes is the type of variation that is available to respond to selection. In hermaphrodites, there is presumably strong selection on alleles that mediate resource allocation to male- vs. female-reproductive success, a within-individual level trade-off that does not occur in diecious species.

Above we noted that stronger local adaptation to males than females observed in the novel thermal regime could be due to novel environments aligning selection between the sexes and selection being, on average, stronger in males than females. In support of the idea that the novel environment aligned selection between the sexes, we found that, within chromosome pools, fitness is more positively correlated between the sexes in the novel than the benign thermal regime (**FIGURE 3**). However, there are, in principle, two possible explanations for this result. One is the aforementioned idea that novel environments re-align sex-specific selection, i.e., GxE for fitness. Alternatively, differences in allele frequencies in the chromosome pools resulting from their different histories of selection in different thermal regimes may be responsible. Prior evidence supporting the idea that *r_MF_* should be *more* positive in novel compared to well-adapted environments has been found in a few studies (Han and Dingemanse, 2017; Berger et al., 2014). While the majority of estimates of *r_MF_* in novel environments have been positive, estimates of *r_MF_* in well-adapted environments are more mixed (Martinossi-Allibert et al., 2018; Koch et al., 2020; Punzalan et al., 2014; Delcourt et al., 2009; Chippendale et al., 2001; Foerster et al., 2007) and, in general, the power to detect differences in *r_MF_*between environments is often low. While we cannot distinguish if the result in our study is due to being assayed in different thermal regimes or due to how selection re-shapes variation within those thermal regimes, it suggests that novel environments do align selection between the sexes more than in benign or better adapted environments.

### Mate Harm

Mate harm may be mediated by traits expressed in both sexes such that the benefit to male fitness of an allele that causes mate harm can be offset not only in the reduced fitness of the male’s female mating partner (i.e., conflict over mating interaction), but also in the reduced fitness of his female offspring (i.e., conflict over shared genetics). If so, then separating the gene pool may result in divergence in traits mediating male harm even though our manipulation of inheritance does not directly affect the conflict over mating interaction. Consistent with previous literature on male-limited selection, we found that male-limited selection led to greater mate harm (Melo-Gavin et al., 2025; Martinossi-Allibert et al., 2019; Rice, 1996; Rice, 1998; but see Jiang et al., 2011). Females housed with ML-males (compared to FL-males) showed a reduction in two measures of fitness: number of offspring per surviving female (productivity) at both temperatures and survival in the novel thermal regime (**FIGURE 4**). This suggests that, under normal inheritance, there may be sex differences in selection for alleles that affect mate harm.

Though the observed divergence in mate harm could be due to increased harmfulness of the ML pool or decreased harmfulness of the FL pool, either scenario likely reflects a response to sex-specific selection pressures as 15 generations is unlikely to be enough time for drift or relaxed selection on male-limited traits in the FL pool to cause such large changes.

Mate harm is thought to evolve as an indirect by-product of competition among males for mating and fertilization success, i.e., the traits responsible for mate harm improve male competitive fitness (Parker, 1979). We see two levels of evidence—among and within chromosome pools—that increased mate harm is associated with elevated male fitness. First, ML-males are both more harmful and more fit than FL-males. Within chromosome pools, we found a negative correlation between male relative competitive fitness and the number of offspring produced by females with whom they were housed (**FIGURE 4**). In principle, the observed differences in the number of offspring per female could be due to differences among males in traits unrelated to mate harm, such as differences in male fertility or larval viability.

However, under either of those alternatives, female productivity would be positively correlated with male competitive fitness, but we find the opposite.

Females housed with ML-males had decreased survival (compared to when housed with FL-males) only in the novel thermal regime (assayed at 28°C), and not in the benign temperature (25°C), suggesting not only increased harmfulness of novel temperature ML-males compared to FL-males, but perhaps also compared to benign temperature ML-males. Because each chromosome pool was only assayed in its own respective thermal regime, we cannot disentangle whether the observed increase in mate harm is due to being assayed at a high temperature or is a product of adapting to high temperature. That is, benign temperature ML-males could be just as harmful as novel temperature ML-males if assayed in the novel thermal regime. However, previous studies have found that assaying males at a higher, stressful temperature results in a reduction mate harm (Martinossi-Allibert et al., 2019; Londoño-Nieto et al., 2023; García-Roa et al., 2018; Londoño-Nieto et al., 2025), suggesting the assay temperature is unlikely to cause the direction of difference that we observed.

Alternatively, adaptation to higher temperature has been shown to produce different effects. Londoño-Nieto et al. (2025) examined mate harm in populations of *Drosophila melanogaster* that had spent ∼29 generation adapting to either 24°C or 28°C. Lines adapted to 28°C were more harmful when assayed at 24°C than when assayed at 28°C (i.e., being assayed at higher temperature lowered the amount of mate harm) *and* lines adapted to 28°C were more harmful at 24°C than lines adapted to 24°C, suggesting that adaptation to higher temperature promoted increased harmfulness. This suggests that adaptation to high temperatures specifically might favour alleles that increase mate harm (such as by favouring alleles that increase sperm protein stability and thus increase the harmfulness of toxic seminal fluids; Martinet et al., 2023; Londoño-Nieto et al., 2025). Alternatively, male-limited selection may alter this plastic decrease in mate harm at high temperatures. Martinossi-Allibert et al. (2019) examined mate harm in seed beetles subjected to male-limited selection for 16-20 generations and found that heat stress lowered the amount of mate harm in populations adapting under normal inheritance but did not change the harmfulness of males in populations adapted to a male-limited selection regime.

Taken together, despite our results that suggest sex-specific selection is more aligned in novel environments (i.e., less intralocus sexual conflict), there may still be selection for increase mate harm regardless.

## Conclusion

We provide evidence that sex-specific adaptation may be hindered by the shared genome, and the effect of the shared genome differs across sexes and across environments. There is growing interest in examining how the sexes adapt to environments differently (Svensson et al., 2018) but understanding how sharing a genome affects adaptations in each sex has been relatively underexplored. Symmetrical sex-limited experimental evolution in different environments is one useful tool to gain insight into these differences.

## Author Contributions

C.M.G. conducted experimental evolution, experiments and analysis and prepared manuscript. M.J.L conducted experimental evolution. A.F.A. designed the experimental evolution, advised on analysis and prepared manuscript.

## Acknowledgements

We are grateful for Yichen Wei, Jeremy Chan, Jamie Chew, Grace Elman, and everyone who helped us maintain these populations.

