## Supplemental Material for "Local Adaptation to the Sex Environment: Reciprocal Sex-Limited Selection in Different Thermal Regimes"

Supplement

*Supplemental methods:*

**History of source population**

*Overview:* The source population used for experimental evolution was initially created by mixing more than 25 inbred lines (from multiple sources) and allowing them to recombine for many generations to generate a diverse set of haplotypes. In this process, two recessive phenotypic markers on Chromosome 3, $se^1$ and $hh^{bar3}$, were introgressed into the population as these were useful for sorting flies during experimental evolution. The end result of this multi-year preliminary phase were two populations. One of these—called the "Grand Ancestor" population—contained genetically diverse copies of Chromosome 3 carrying $se^1$. The other population—called the "Ancestor" population—was created from a copy of the Grand Ancestor and contained genetically diverse copies of Chromosome 3 carrying $se^1$ and $hh^{bar3}$. The other chromosomes (Chromosomes 2, 4, *X* and *Y*) were also genetically variable in both of these populations but are not the focus here. Below is a more detailed description of how this source population for our experimental population was created. Some of the steps in the creation of these populations appear capricious in retrospect as plans changed over the multi-year period prior to the onset of experimental evolution but all the steps are reported below for completeness.

First, we mixed 23 lines from the Drosophila Genomic Resource Panel (DGRP; Mackay et al., 2012) and 3 African lines (received from John Pool's lab at the University of Wisconsin) as follows. In April 2019, two subpopulations were created. First, DGRP177 and DGRP507 and 3 in-bred African lines (SP159N, EF81N, and EF122N) were each crossed to an inbred line homozygous for *sepia* mutant allele ($se^1$). *Sepia* is a recessive eye-colour marker located in position 66D on Chromosome 3R (Öztürk-Çolak et al., 2024). F1s from each cross were then crossed to two other F1 lines and subsequent F2s were combined to create this first subpopulation. Concurrently, a second subpopulation was created where 10 other DGRP lines (DGRP42, DGRP177, DGRP235, DGRP427, DGRP439, DGRP508, DGRP703, DGRP765, DGRP799) were also crossed to the $se^1$ mutant line, and F1s were crossed to two other F1 lines. After 18 generations of maintenance of these subpopulations, a third subpopulation was created from crossing males from subpopulation 2 with virgin females from 11 more DGRP lines (DGRP57, DGRP195, DGRP208, DGRP315, DGRP357, DGRP371, DGRP391, DGRP395, DGRP399, DGRP491, DGRP509, DGRP517, DGRP516, DGRP843).

These three subpopulations (each segregating for $se^1$) were maintained separately until April 2021, when the three subpopulations were equally mixed to create one population we refer to as the "Grand Ancestor." Once mixed, the Grand Ancestor population was kept at a population size of $N = 1600$ corresponding to 200 individuals per bottle (40mL standard yeast media) and kept at 25° C on a two-week cycle at 50% relative humidity. One additional bottle of 200 $se^1/se^1$ homozygotes flies was maintained separately. Each generation, this subpopulation was composed of 180 $se^1/se^1$ individuals from the previous generation's $se^1/se^1$ bottle as well as 20 $se^1/se^1$ individuals from the main populations. Reciprocally, 20 individuals from the $se^1$ subpopulation were mixed into the main population each generation. This Grand Ancestor population was maintained this way for ~50 generations (totalling > 90 total generations of lab adaptation).

Then we introgressed the $hh^{bar3}$ marker into a copy of the Grand Ancestor population to create the "Ancestor" population. The stock containing $hh^{bar3}$ was backcrossed to the Grand Ancestor population for ~50 generations prior using the two-generation cycle illustrated in **FIGURE S3**. This population contained $hh^{bar3}$ at a high frequency but was genetically variable for the rest of the genome. After this backcrossing, the *hh*$^{bar3}$ stock was expanded and crossed to the Grand Ancestors to produce a population that was homozygous for genetically diverse copies of both *se*$^{1}$ and *hh*$^{bar3}$. This occurred as follows. 210 females homozygous for *hh*$^{bar3}$ were crossed to 205 *se*$^{1}$/*se*$^{1}$ Grand Ancestor males in 10 vials that were flipped into new vials twice (resulting in 30 vials) to generate a large number of offspring. Approximately 1700 F1s were mated amongst themselves (in ~30 vials). The following generation we collected 700 *hh*$^{bar3}$/*hh*$^{bar3}$ females (irrespective of *se*$^{1}$ genotype) and crossed them to 713 *se*$^{1}$/*se*$^{1}$ Grand Ancestor males. Once again, the resulting offspring mated amongst themselves. From the following generation, ~1500 flies homozygous for both *hh*$^{bar3}$ and *se*$^{1}$ were collected and mated to one another (in 10 bottles with ~150 flies per bottle) to establish the "Ancestor" population, which was maintained on a two-week cycle at 25° C with 50% relative humidity). This Ancestor population was homozygous for both *hh*$^{bar3}$ and *se*$^{1}$ but should otherwise be genetically variable (and equivalent to the "Grand Ancestor" population, which lacks *hh*$^{bar3}$).

The Ancestor population was maintained this way for 8 generations and then an additional pulse of variation was introduced by adding 170 *se*$^{1}$/*se*$^{1}$ Grand Ancestor males into this Ancestor population (which were otherwise *se*$^{1}$ *hh*$^{bar33}$). ~2200 (unsorted) offspring were then used to establish the next generation (distributed across 15 bottles). The next generation the Ancestor population returned to the standard maintenance schedule ($N$ = 1500, *se*$^{1}$ *hh*$^{bar3}$ individuals as described above.).

Two *TM6B* balancer stocks were also used to establish the experimental populations to aid with fly sorting. One of these *TM6B* balancers was marked with the dominant phenotypic bristle marker *Sb* and contained a construct with a dominant fluorescent marker, *DsRed* (the full genotype of the construct, *DsRed Hr5.IE1 OpIE2-neoR traF OpIE2-pacR dsx* , is described in

Kandul et al., (2019)). We hereafter referred to this balancer as *TM6B DsRed Sb.* The other balancer was only marked with the *Sb* marker and is referred to as *TM6B Sb. TM6B Sb* and *TM6B DsRed Sb* are recessive lethal allowing us to track their inheritance each generation and thereby infer inheritance of focal chromosomes as per our experimental evolution regime. Both balancer stocks were backcrossed 2-4 times to the Ancestor prior to their use with each backcross generation using hundreds of Ancestor flies. The final stock population for each balancer involved ~1200 flies from the Ancestor. (Note, the non-balancer 3$^{rd}$ chromosomes in these two crosses were exclusively from the Ancestor population).

From these two stock populations, ~1000 *TM6B DsRed Sb/*$se^1$ $hh^{bar3}$ females were collected and mated to ~1000 *TM6B Sb/*$se^1$ $hh^{bar3}$ males in 13 bottles (40mL standard yeast media, roughly 150 flies per bottle) to begin Generation 0 of the experimental populations. This cross was flipped into new bottles 5 times (over 12 days), resulting in 6 sets of 13 bottles, one for each of the 6 experimental populations. Three of these were placed at 25° C for the benign temperature treatment and the other three at 28° C for the novel temperature. The offspring from this cross were considered Generation 1 and were sorted following our experimental design.

Due to unforeseen circumstances, one population (the third replicate population for the benign temperature), failed to provide sufficient number of individuals to support generation 1 and was restarted using unused offspring from the other five populations.

For each population experimental sorting proceeded as follows. Each generation, on Day 14, flies were mixed in small cages and sampled by aspiration. After sorting by sex, flies were briefly stored in sex-specific vials (N=150 flies per vial, 10 vials of each sex). Within the next three hours, $hh^{bar3}$ $se^1$ homozygous individuals were removed from the population and then males and females were sorted for presence/absence of the fluorescent *DsRed* marker to distinguish which of the *TM6B* balancers they had inherited and, therefore, which of the focal chromosomes—Male-Limited or Female-Limited—they had inherited. For example, in generation one, fluorescent individuals must have inherited *TM6B DsRed Sb* from their mothers and so must have inherited their focal chromosome from their fathers.

Each generation, 520 males and 520 females were selected such that they had inherited their focal Chromosome 3 from their same sex parent and were placed in new vials (7 mL standard media) in a ratio of 13 males: 13 females and maintained at one of two treatment temperatures (25° C for the benign temperature populations and 28° C for the novel temperature populations). Flies interacted for five days before being transferred into new vials (7 mL standard media). (This transfer was done to ensure quality of the food did not deteriorate and cause stress on the flies during the interaction stage; the second set of interaction vials also served as "back-up" vials for offspring production.) After a total of 7 days of male/female interaction, flies were transferred into oviposition vials with fresh yeast (7 mL standard media) and allowed to oviposit for 24-28 hours after which adults were discarded, thereby initiating Day 1 of the next generation. For the novel temperature treatment, after oviposition, vials were held for 2 days at 18° C and then returned to 28° C. This was done to ensure synchronization of generations between the benign and novel temperature treatments (i.e., all populations on the same three-week cycle). Then on Day 14 of the new cycle, flies were mixed in small cages and sorted again.

In generations where the main oviposition vials provided insufficient number of offspring of the correct genotypes, additional offspring from the "back-up" vials were used to supplement the main populations. Back-up flies were sampled and sorted the same way as described above. For each temperature treatment, there was one generation with unforeseen logistical challenges

(Generation 10 for the benign temperature and Generation 2 for novel temperature) that required us using back-up vials for 50% of flies that generation. Otherwise, back-up flies accounted for, on average ~7% of benign temperature treatment flies, and less than 2.9% of novel temperature treatment flies.

*Supplemental Figures*

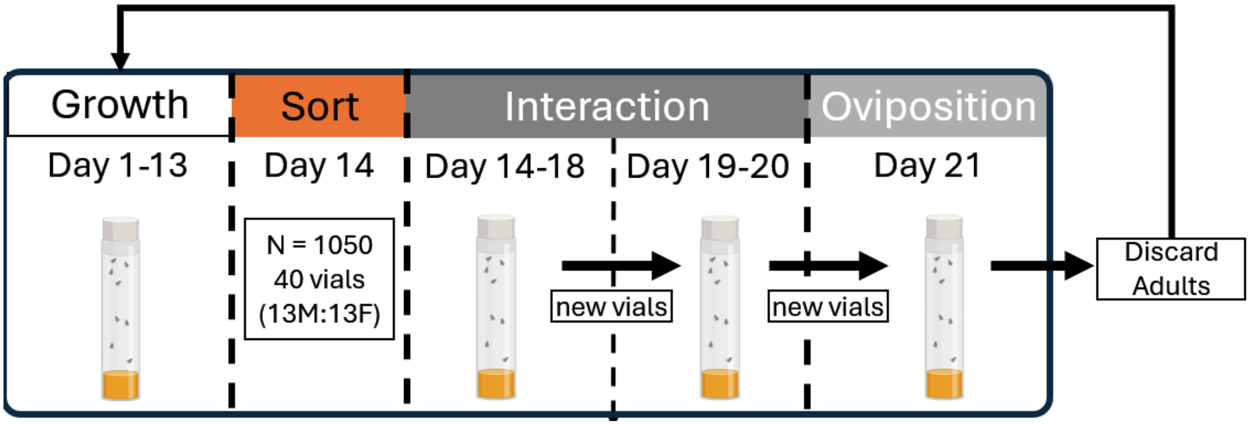


**SUPPLEMENTAL FIGURE 1**: Three-week maintenance protocol for experimental populations. All 6 populations were maintained on a three-week cycle (12:12 hour light:dark cycle at 50% relative humidity) on regular yeast-sugar media. Flies were mixed in small cages on Day 14 of each generation and then sorted (as described in **FIGURE 1**) into interaction vials (40 vials of 13 males paired with 13 females for each population). After seven days of interaction, flies were transferred to oviposition vials to lay eggs for 24-28 hours, after which adults were removed, beginning Day 1 of the next generation cycle.

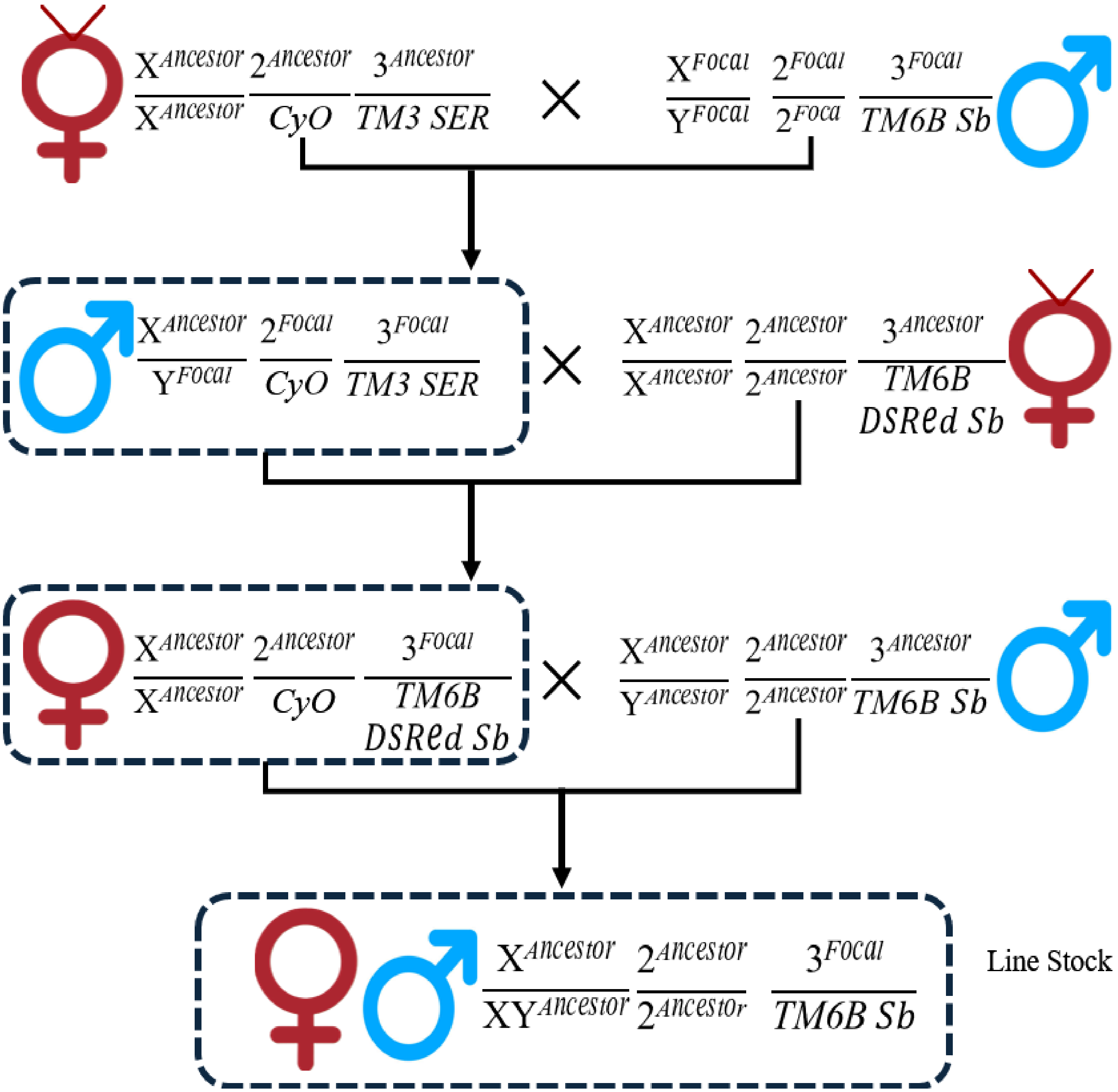


**SUPPLEMENTAL FIGURE 2** Isolation of ML and FL chromosomes into lines onto the Ancestral Background. Approximately 40 individual male flies from each chromosome pool (FL or ML) for each population were obtained at generation 15 and then went through the following series of crosses to isolate their focal chromosome onto the Ancestral Background. The "V" symbol above the female symbol indicates when virgin females were used. Except for the focal male in the first cross that was isolated individually, 3-5 individuals of each sex were used for each subsequent cross.

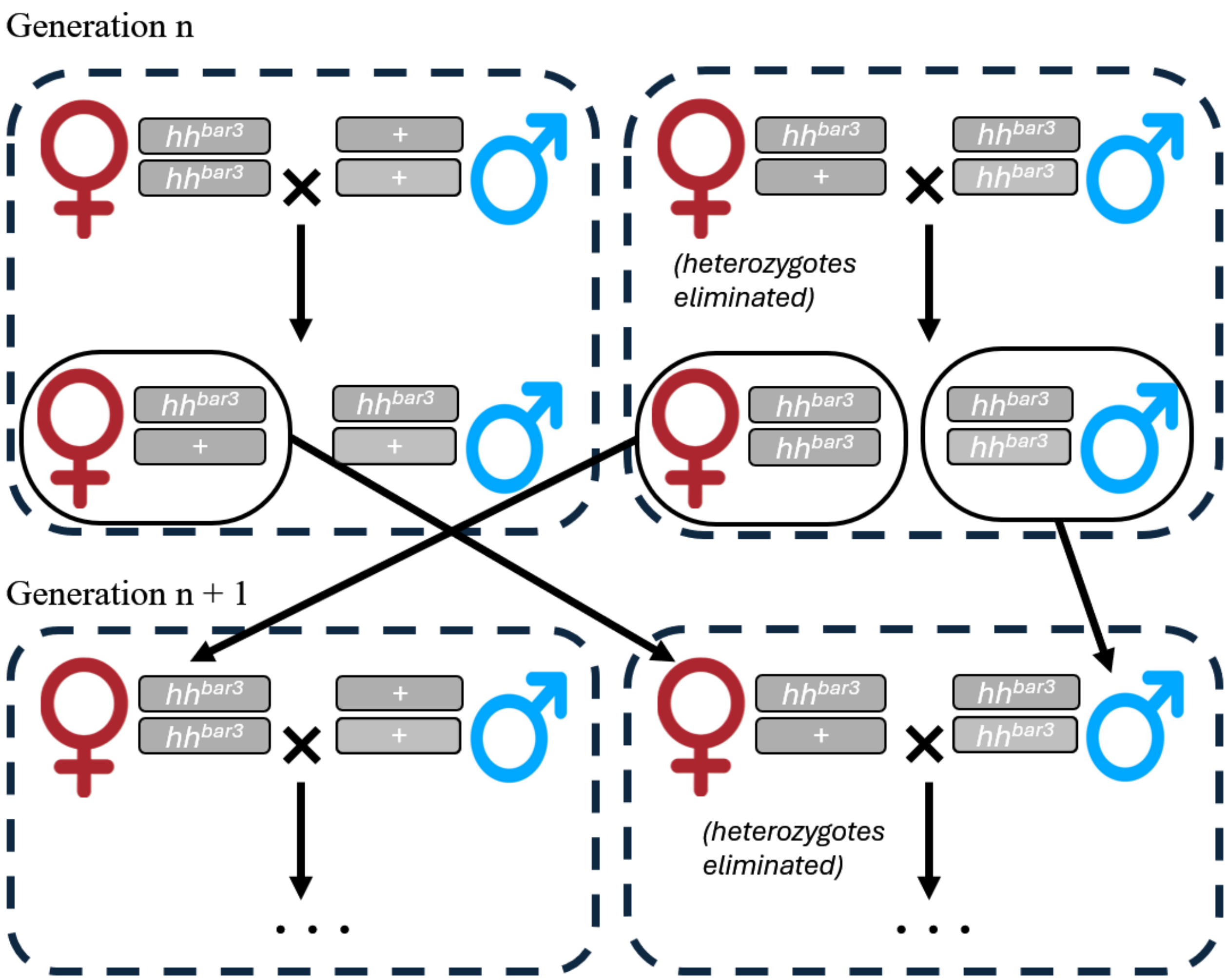


**SUPPLEMENTAL FIGURE 3:** Backcross procedure of the marker $hh^{bar3}$ into the Grand Ancestor Population. Each cross contains 3 vials of N ≈ 100. In generation *n*, $hh^{bar3}$ females from generation *n* - 1 are crossed to Grand Ancestor males (left). Simultaneously, $hh^{bar3}$ /+ females are mated to $hh^{bar3}$ *males* (right). Then, in generation n + 1, F1 hybrid females (derived from left cross) are mates to homozygous $hh^{bar3}$ males (derived from the right cross). Simultaneously, $hh^{bar3}$ females from generation *n* are crossed to Grand Ancestor males (left).

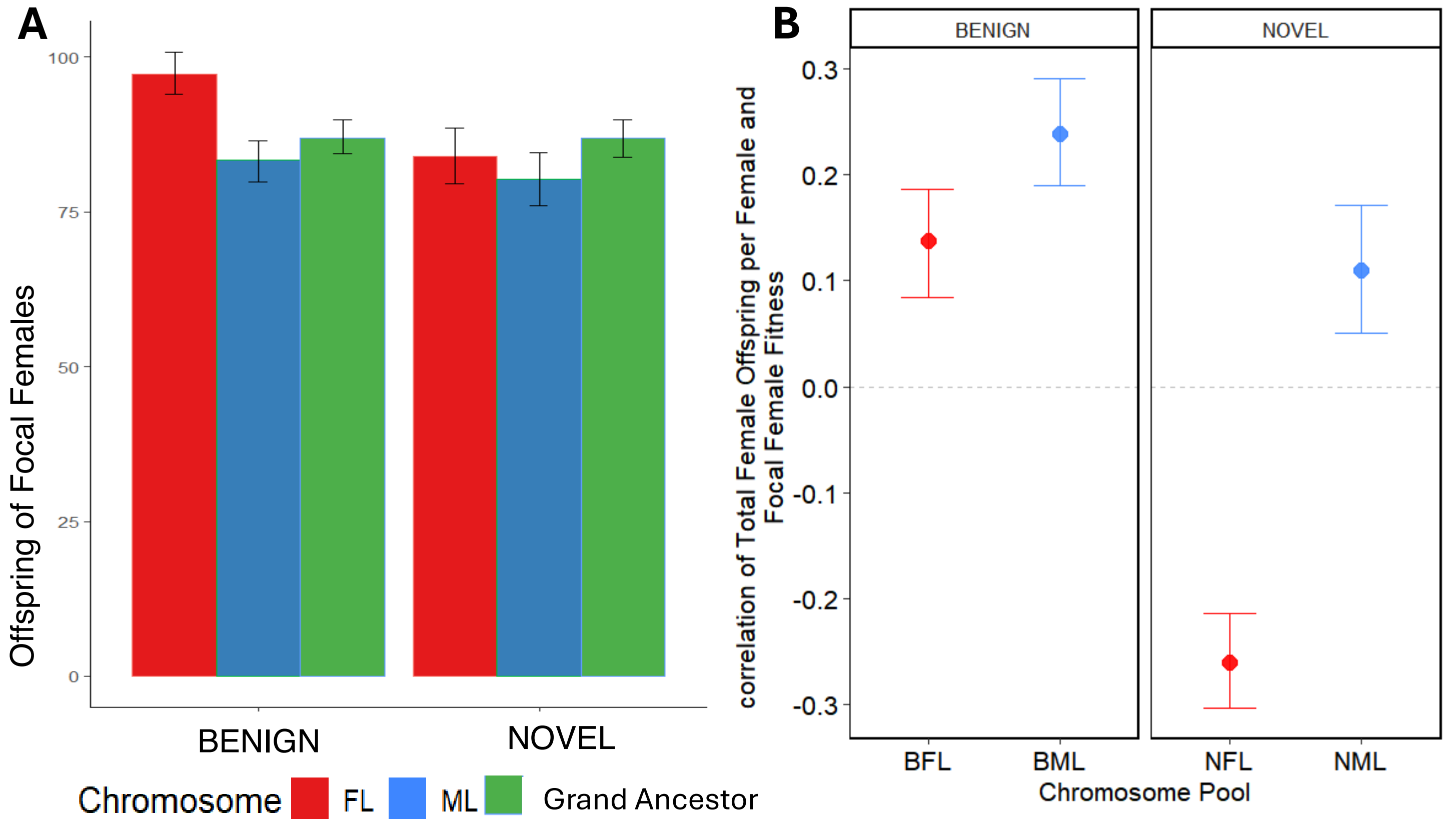


**SUPPLEMENTAL FIGURE 4**: Focal female fecundity when carrying ML or FL chromosomes. (A) Number of offspring produced by focal females in the female competitive fitness assay. Error bars are 95% confidence intervals of the means. (B) The Pearson correlation between fecundity of focal females (the number of offspring produced by focal females) and the competitive fitness of females. We centered fecundity (*f* = number of focal offspring produced per replicate vial) for each line with respect to the mean fecundity for each chromosome type (*C*) per each population (*P*) $\tilde{f}_{S,C,P,i} = f_{S,C,P,i} - \bar{f}_{S,C,P}$. Competitive fitness was also centered. The Pearson correlation between centered female competitive fitness and centered female fecundity was calculated for each chromosome type within each temperature treatment (novel temperature ML, novel temperature FL, benign temperature ML and benign temperature FL). We performed 1000 bootstraps for each chromosome type by resampling lines with their centered female fitness and fecundity values and calculating the Pearson correlation coefficient for each sample. Unexpectedly, FL-females in the novel thermal regime showed superior competitive fitness (compared to ML-females; Figure 2B) without any difference in *number* of focal offspring produced (shown here). This is in contrast to benign temperature FL-females, which had higher competitive fitness than ML-females (Figure 2B) and produced considerably more focal offspring (shown here). This result at novel temperature shows that total female fitness may be greatly affected by fitness components other than those directly affecting number of offspring (productivity) *per se*, such as female-female competition concerning acquisition of food resources or over ideal oviposition sites or oviposition timing. Indeed, when examining variation within pools, competitive female fitness is positively correlated with the number of focal offspring per female in all chromosome pools except FL in the novel thermal regime. Pearson's correlation: $r_{BFL}$ = 0.138 [0.190, 0.0881], $r_{BML}$ = 0.238 [0.288, 0.186], $r_{NML}$ = 0.107 [0.175, 0.0510], $r_{NFL}$ = -0.260 [-0.220, -0.302]).
